# Testing of various protocols for human skin cryopreservation by vitrification method

**DOI:** 10.1101/2023.11.22.568219

**Authors:** Andrei Riabinin, Maria Pankratova, Olga Rogovaya, Ekaterina Vorotelyak, Andrey Vasiliev

**Affiliations:** Koltzov Institute of Developmental Biology of Russian Academy of Sciences, 26 Vavilov Street, Moscow, Russia, 119334; Moscow States University, Biological Faculty, 1-12 Leninskie Gory, Moscow, Russia, 119234

**Keywords:** skin, vitrification, MTT, DMSO, glycerol, sucrose, polyethylene glycol

## Abstract

Optimization of skin cryopreservation technology is an important task for translational medicine, pharmacology, and a list of other fields. One of the promising approaches is vitrification. This rapid freezing technique allows one to ensure minimal damage to the sample by equating the formation of ice crystals and dehydration both in the cytoplasm of cells and in the extracellular space. This study is aimed at finding the optimal combination of cryoprotective agents (DMSO, glycerol, sucrose and polyethylene glycol) to achieve the highest viability of skin samples after vitrification.

## Introduction

While cell cryostorage has been developed long ago, preservation of tissues and their engineered equivalents is still challenging (Hunt, 2017). One of the important aspects of successful cryopreservation is the mode of temperature decline. The two main approaches in this respect are systemic slow freezing and vitrification, very fast freezing (Fahy and Wowk, 2015; Whaley et al., 2021). Vitrification involves instant freezing when the liquid in the sample passes to an amorphous glassy state with a transition to the amorphous ice phase at -130 ° C evading the crystallization phase with the formation of symmetrical hexagonal crystals (Fahy and Wowk, 2015; Hunt, 2017).

Optimization of cryopreservation methods is aimed at increasing cell viability after full cycle of cryopreservation, leveling defects in extracellular structures of tissues and organs and reducing the cost of the process itself (Chen et al., 2023). The main ways to improve cryopreservation technology are the choice of optimal temperature programs for freezing, storage and defrosting, the search for new cryoprotective agents (CPA) and methods of tissue processing that would allow better resistance of the samples to the negative temperature’s effects. Due to their relatively thinness and relatively low density, human skin and its components, such as epithelium, are a common biological object, for which various cryopreservation techniques can be effectively applied, and many cryopreservation protocols have been developed. In this work, several types of cryopreservants were tested, which included combinations of CPA often used for skin vitrification (Tian et al., 2019; Zhu, 1991; Zieger et al., 1997; Son et al., 2021).

Extracellular CPA are compounds with a high molecular weight that are unable to penetrate the cell membrane thus having less cytotoxic effect, and they are much easier to remove from the tissue after defrosting (Linkova et al., 2022). The mode of action of non-penetrating CPA is based on binding extracellular water molecules preventing the growth of ice crystals. This protects cells from osmotic changes and allows reduction of more cytotoxic CPA (such as dimethylsulphoxide, DMSO) concentrations (Whaley et al., 2021).

Polyethylene glycol (PEG), which is an extracellular CPA with undetermined potential, was also tested in this research. It had not previously been used for cryopreservation of the skin but had performed well in cryopreservation of other objects (Oltean et al., 2012; Lee et al., 2013; Puts et al., 2015).

## Materials and methods

### Samples

Skin samples were obtained during plastic surgery procedures at the Medical Research Center of High Medical Technologies – Central Military Clinical Hospital named after A.A. Vishnevsky of the Ministry of Defense of the Russian Federation in accordance with the Agreement on scientific cooperation between this center and N.K. Koltzov Institute of Developmental Biology of the Russian Academy of Sciences (IDB RAS) with the informed consent of the patients.

Skin fragments and epidermal layers from 3 different donors were used in the experiments (N=3). For each experimental group, 5 technical repetitions were performed.

### Processing of donor material

After receiving the skin, it was washed in the sufficient volumes of Hanks solution (PanEco) with an antibiotic (gentamicin (PanEco)). Then the hypodermis was cut off using a scalpel, and the flap of skin was cut into equal round fragments with a diameter of 5 mm using a scalpel.

To obtain skin sheets, some of the resulted skin fragments were incubated for 24 hours at +4 °C in 0.2% dispase (Sigma). After that, an epithelial layer was mechanically separated from the dermis using forceps.

### Skin cryopreservation

5x5 mm skin layers and fragments were incubated for 30 minutes in the cryopreservation medium (Tables 1-- 7) in sterile retort bags at room temperature. Three skin fragments and three epithelial layers were placed in each bag. After incubation, the bags were sealed, placed in liquid nitrogen and transferred to the cryopreservation room.

**Table 1.** Cryopreservation medium 1: DMSO + fetal bovine serum (FBS)

| Component | Manufacturer | Concentration |
| --- | --- | --- |
| RPMI 1640 | PanEco | 1x |
| DMSO | Sigma | 20% |
| FBS | HiMedia | 10% |
| EGF | Sigma | 10 ng/ml |

**Table 2.** Cryopreservation medium 2: DMSO + FBS + sucrose.

| Component | Manufacturer | Concentration |
| --- | --- | --- |
| RPMI 1640 | PanEco | 1x |
| DMSO | Sigma | 20% |
| FBS | HiMedia | 10% |
| Sucrose | Sigma | 0.5M |
| EGF | Sigma | 10 ng/ml |

**Table 3.** Cryopreservation medium 3: Glycerol + FBS.

| Component | Manufacturer | Concentration |
| --- | --- | --- |
| RPMI 1640 | PanEco | 1x |
| Glycerol | Sigma | 20% |
| FBS | HiMedia | 10% |
| EGF | Sigma | 10 ng/ml |

**Table 4.**
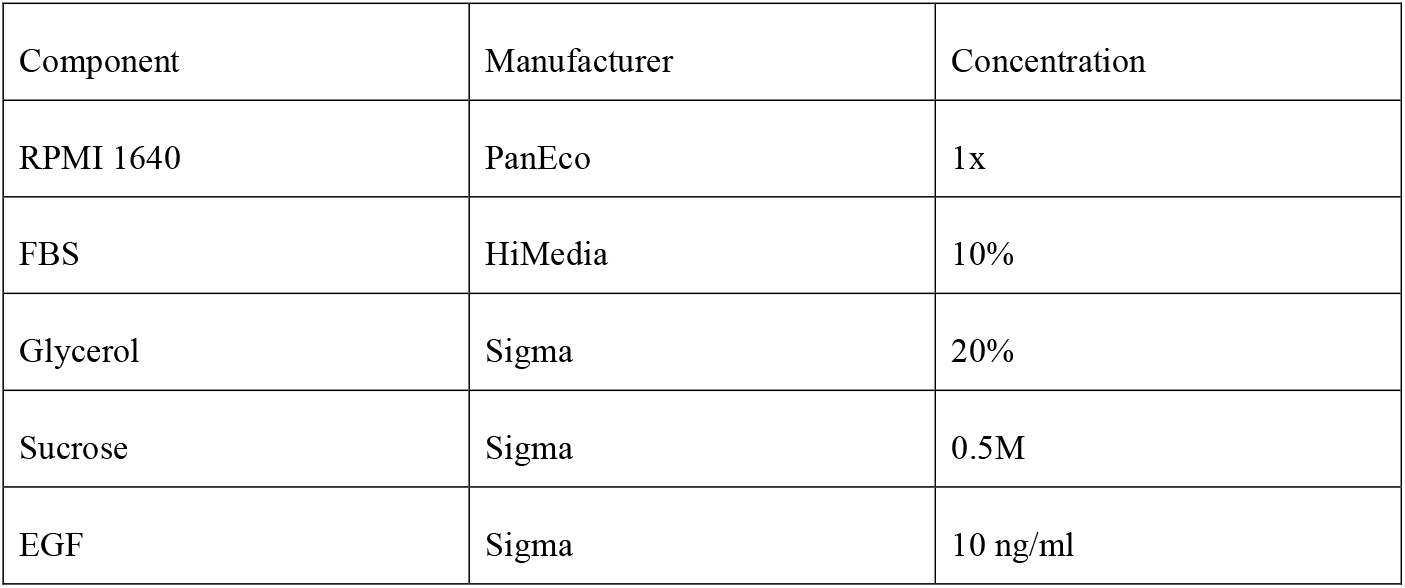
Cryopreservation medium 4: Glycerol + FBS + sucrose.

| Component | Manufacturer | Concentration |
| --- | --- | --- |
| RPMI 1640 | PanEco | 1x |
| FBS | HiMedia | 10% |
| Glycerol | Sigma | 20% |
| Sucrose | Sigma | 0.5M |
| EGF | Sigma | 10 ng/ml |

**Table 5.** Cryopreservation medium 5: DMSO.

| Component | Manufacturer | Concentration |
| --- | --- | --- |
| RPMI 1640 | PanEco | 1x |
| DMSO | Sigma | 20% |
| EGF | Sigma | 10 ng/ml |

**Table 6.** Cryopreservation medium 6: DMSO + PEG+EGF.

| Component | Manufacturer | Concentration |
| --- | --- | --- |
| RPMI 1640 | PanEco | 1x |
| DMSO | Sigma | 20% |
| PEG 6000 | Sigma | 20% |
| EGF | Sigma | 10 ng/ml |

**Table 7.** Cryopreservation medium 7: DMSO + PEG + sucrose+EGF.

| Component | Manufacturer | Concentration |
| --- | --- | --- |
| RPMI 1640 | PanEco | 1x |
| DMSO | Sigma | 20% |
| PEG 6000 | Sigma | 20% |
| Sucrose | Sigma | 0.5M |
| EGF | Sigma | 10 ng/ml |

### Thawing and viability assessment of fresh and cryopreserved samples

Thawing of bags with samples was carried out a week after cryopreservation by immersion in a water bath at +37 °C.

Thawed skin fragments were transferred to the wells of a 48-well plate. After this, 300 μl of medium RPMI with 10% MTT was poured into the wells. The plate with samples was incubated for 4 hours in a CO_2_-incubator at +37°C on an orbital shaker at 65 rpm. After this, the medium with MTT was taken and 300 μl of DMSO was added. Dissolution of tetrazolium salts in DMSO was carried out on a 3d shaker at 60 rpm for 12 hours. After the appropriate incubation time, 100 μl of the solution from each well of a 48-well plate with samples was transferred to the wells of a 96-well plate and analyzed on a BioTek Synergy H1 spectrophotometer at a wavelength of 545 nm.

Negative control: skin fragments treated with a 10% solution of Triton-X100 (Sigma) in phosphate buffered saline (PanEco) for 30 minutes at +37 °C in a CO_2_-incubator on an orbital shaker at 65 rpm. Positive control: fresh skin fragments before freezing.

To standardize the results of MTT measurements according to mass, the values measured by the spectrophotometer were divided by the values of the masses of the corresponding skin fragments and epidermal layers.

### Statistical analysis

Statistical analyses in all experiments were performed using software STATISTIKA (v.10.0). ANOVA single factor test was used to compare MTT experiment data between negative control and experimental groups and between cryopreservation medium 7 and other experimental groups. Results with p values < 0.05 were considered significantly different from the null hypothesis. The data on the graphs are presented as an average value ± standard deviation.

## Results and discussion

The effectiveness of several cryoprotectors for vitrification of two types of samples, human skin fragments and human epidermal layers, was tested. This method for tissue cryopreservation was chosen based on several studies, in which the vitrification method was tested on human skin. It can be concluded that this method of freezing is quite effective and demonstrates the same or higher level of sample integrity, morphology preservation and cell viability in comparison with systemic freezing (Campbell and Brockbank, 2022). In addition, this method does not require specific equipment for programmable slow freezing of the skin, which simplifies the protocol itself.

The results obtained when assessing the viability of tissues after defrosting using MTT analysis showed that one of the most effective formulations of cryoprotector for skin vitrification in comparison with other experimental groups is a combination of intracellular CPA DMSO and extracellular CPAs: polyethylene glycol and sucrose, Δ (MTT value) was -17% (Fig. 1). For vitrification of the epidermis, the combination of DMSO, PEG and sucrose was less effective, Δ(MTT value) = -40%, but the viability of thawed samples remained at an acceptable level.

**Fig. 1.**
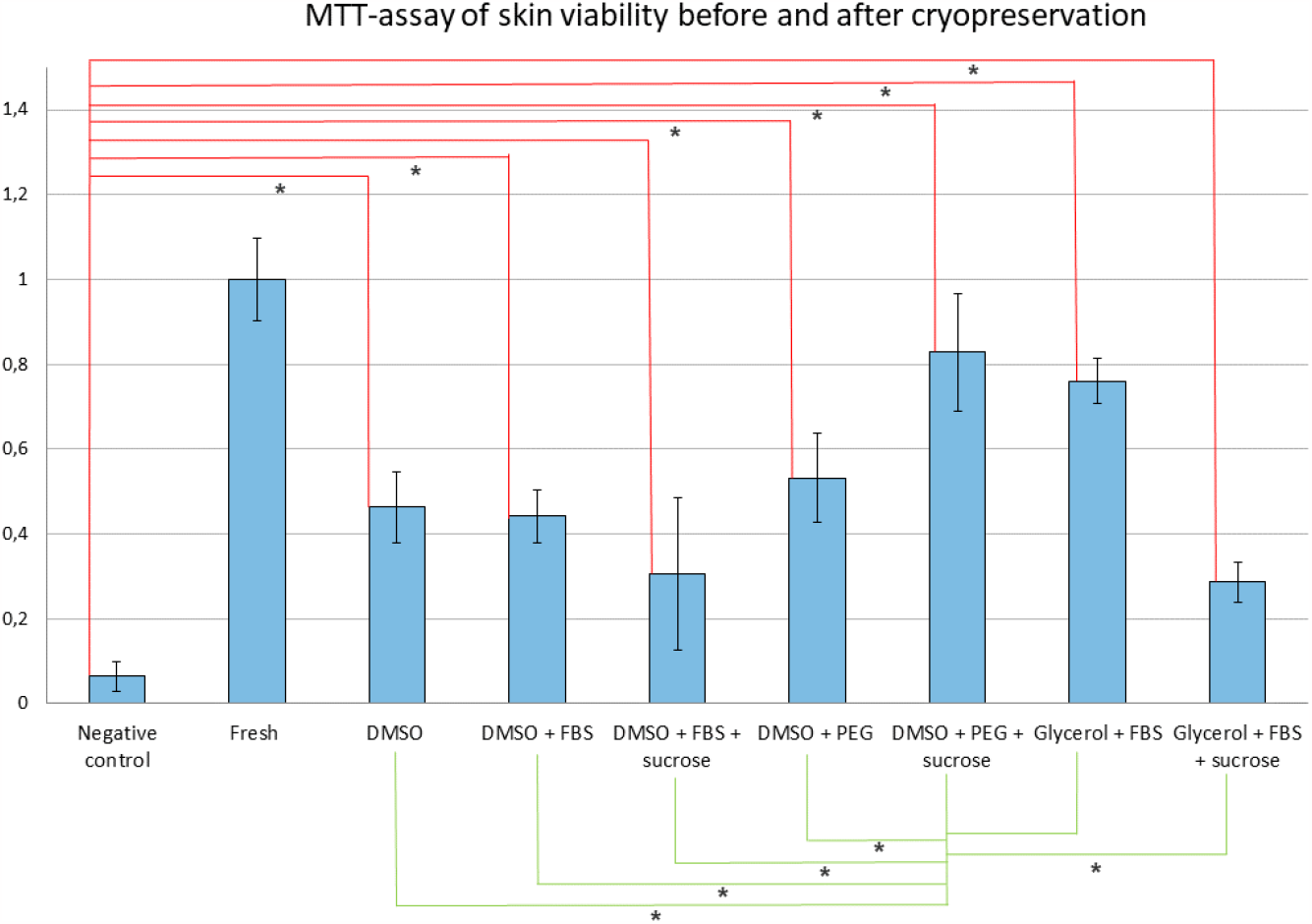
Relative coefficients of the light absorption of formazane solution collected from the wells with negative control, positive control (fresh), and experimental groups skin samples.

The results obtained are generally consistent with the literature data, since a similar combination of extracellular and intracellular CPA had previously been successfully used in several studies of skin vitrification (Silvestre et al., 2002; Son et al., 2021). However, in this work, PEG polymer was used as an extracellular CPA for skin cryopreservation for the first time. On other types of tissues and cells, PEG had previously been shown to have many advantages in the preservation of cells and organs, such as antioxidant capacity, stabilization of membranes and protection from exposure to negative temperatures (Oltean et al., 2012; Lee et al., 2013; Puts et al., 2015). In several other studies, PEG in combination with other substances had also been successfully used for vitrification (O’Neil et al., 1997; Derakhshan et al., 2017; Jung et al., 2021). Sucrose also belongs to non-penetrating CPA and is often used in combination with other substances for skin vitrification (Fujita et al., 2000; Silvestre et al., 2002; Son et al., 2021).

Another effective cryopreservant for vitrification of both skin and epidermis is a combination of glycerol with FBS, Δ(MTT value) was -24% (Fig. 1, Fig. 2). In general, glycerol is rarely used in tissue vitrification protocols and is not used for skin vitrification. It can be assumed that such a high efficiency of this composition is associated with low cytotoxic effect of glycerol (Udoh et al., 2000; Schiozer et al., 2013).

**Fig. 2.**
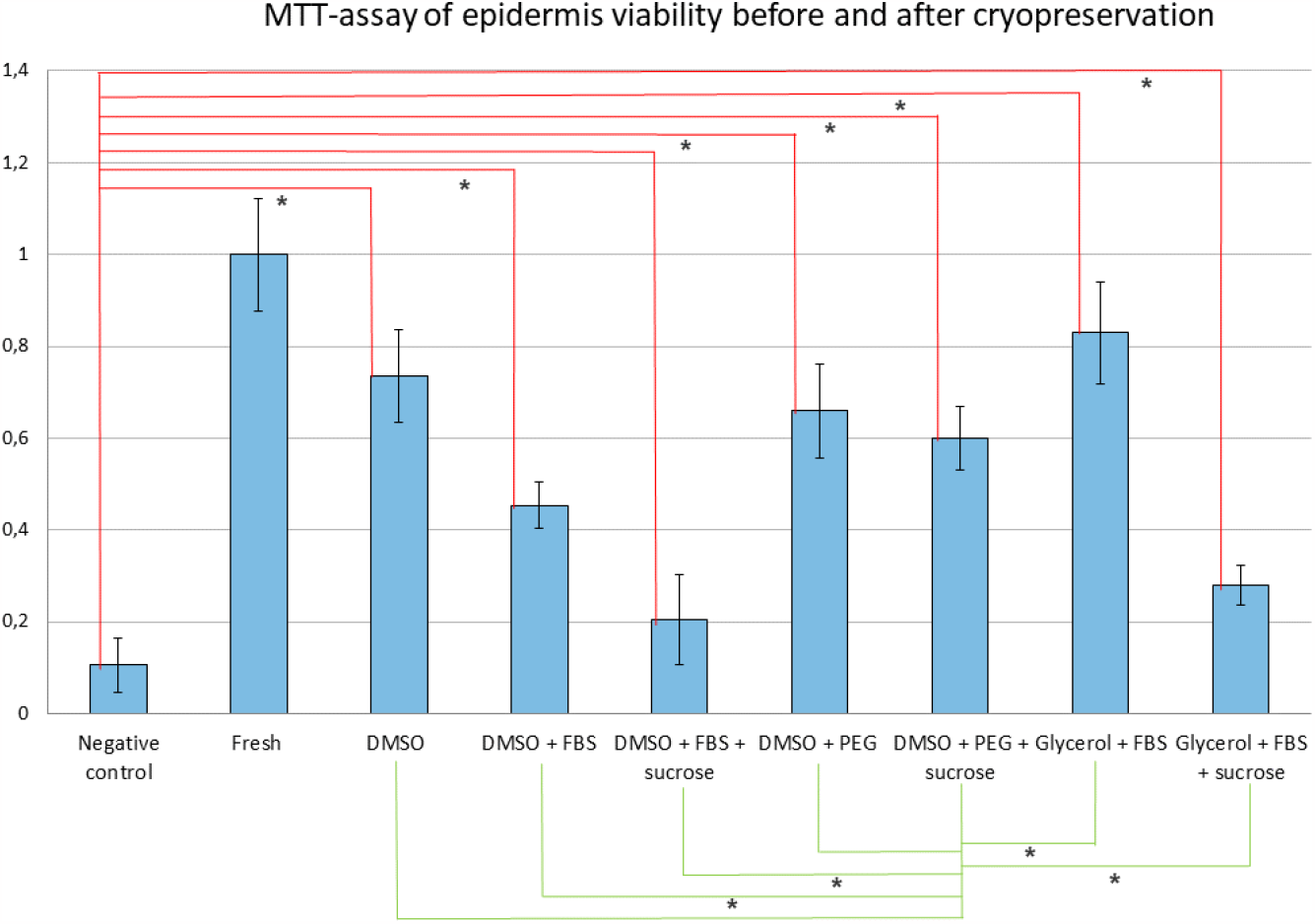
Relative coefficients of the light absorption of formazane solution collected from the wells with negative control, positive control (fresh), and experimental groups epidermal layers samples.

The viability of skin samples and epidermal layers frozen in a cryopreservant made of DMSO, FBS and sucrose (p>0.05) did not differ from the negative control, which allows us to draw conclusions about the ineffectiveness of such a composition (Fig. 1, Fig. 2). In the group where a combination of glycerol, FBS, and sucrose was used, low viability of both skin samples ((Δ(MTT value) = -71%) and epidermis ((Δ(MTT value) = -72%) was observed, also. It is possible that such low efficiency in these groups is associated with cell damage due to high osmotic pressure created in the extracellular space by high concentrations of sucrose and serum components.

It is also worth emphasizing that EGF was added to all tested cryopreservation media as a supporting agent that increases the viability of cells in the sample, at an optimal concentration of 10 ng/ml, according to the literature data (Fahmy et al., 1993). Based on the ability of this protein to prevent cell apoptosis, regulate skin homeostasis, and increase the lifespan of keratinocyte cultures, we suggested that its adding to cryopreservant may increase cell survival (Nanba et al., 2013).

## Conclusions

The resulted data suggest that the developed cryopreservant, which includes 20% DMSO, 20% PEG and 0.5M sucrose, as well as 10 ng/ml EGF (Cryoconservation medium 7), is an effective and promising medium for vitrification of human skin, since it provides higher viability of samples after defrosting.

In the future, within the framework of this topic, it is possible to perform other viability tests, such as the WST1 test, trypan blue staining, the analysis of glucose uptake and lactate production, and the analysis of oxygen consumption. In addition, it is possible to compare the effectiveness of slow freezing and vitrification methods, as well as testing other CPA and their combinations as cryoprotectors.

## Data availability

All data are available upon request to the corresponding author, Andrei Riabinin.

